# Effect of GDF15 on acetaminophen (APAP)-induced liver injury in mice

**DOI:** 10.1101/2022.03.13.484113

**Authors:** Peng Jiang, Zhenghong Liu, Tingyu Fang, Zhidan Zhang, Dongdong Wang, Suowen Xu, Jianping Weng

## Abstract

Acetaminophen (APAP)-induced liver injury (AILI) has been recognized as a pivotal contributor to drug-induced liver failure in western countries, but its molecular mechanism remains poorly understood. After data mining using the Gene Expression Omnibus (GEO) database, we found that the gene expression of growth differentiation factor 15 (GDF15) increased significantly after APAP-induced hepatotoxicity in mouse liver. However, the precise role of GDF15 in AILI has yet to be clarified. To deal with this question, we first examined the temporal gene expression pattern of GDF15 after APAP treatment. Next, overexpression of hepatic GDF15 using adeno-associated virus serotype 8 (AAV8) was employed to investigate the role of GDF15 in AILI. In addition, therapeutic administration of recombinant human GDF15 (rhGDF15) was conducted to probe its curative effect after APAP overdose. We observed that protein expression of GDF15 was significantly increased after APAP overdose. However, there were no significant differences in AILI-related outcomes after overexpression or supplementation of GDF15. The contents of serum alanine aminotransferase (ALT) and aspartate aminotransferase (AST), morphological feature of liver, the level of reduced form of glutathione (GSH), the activation of c-Jun-N-terminal kinase (JNK), the expression of genes relevant to oxidative stress and cell proliferation were not remarkably changed between AAV8-GFP and AAV8-Gdf15 groups. Although the liver injury was not improved by rhGDF15 administration, the gene expression levels of glutamate-cysteine ligase modifier subunit (Gclm) and glutamate-cysteine ligase catalytic subunit (Gclc) which function downstream of Nrf2 signaling pathway were significantly increased. Altogether, despite a noticeable elevated expression after APAP overdose, GDF15 overexpression or supplementation exerted no significant ameliorative effect on AILI. Further mechanism and consequence of Gclm and Gclc elevation by GDF15 remains to be elucidated.

## 1. Introduction

Acetaminophen (APAP) is an antipyretic analgesic with extensive use in clinic. It is usually safe for use at therapeutic doses. However, severe liver injury can occur in patients with overdose of APAP. Recent studies have shown that therapeutic dosage of APAP can cause hepatotoxicity in patients under fasting or excess alcohol drinking (Louvet et al., 2021). APAP-induced liver injury (AILI) is thought to the primary cause of liver failure in western nations (Bernal et al., 2010;Lee, 2017), and it is the most studied event of drug-induced liver damage. The mechanisms of AILI are related with diverse processes such as depletion of glutathione, production of toxic metabolites, oxidative stress, sterile inflammation, autophagy and liver regeneration (Yan et al., 2018). Despite intensive research interest, the precise mechanism of AILI remains largely unclear. Moreover, the only therapeutic option of AILI is n-acetylcysteine (NAC) which has a rather limited therapeutic window of 8 hours (Lee et al., 2009). The development of novel therapeutic drugs for AILI remains an unmet medical need.

With the development of high-throughput sequencing technology, data mining using public databases (including GEO database) makes it possible to elucidate the mechanism of drug action or toxicity. After analyzing the differentially expressed genes (DEGs) from three datasets related to AILI, we found a significant elevation of growth differentiation factor 15 (GDF15) in all the three datasets. Therefore, we supposed GDF15 may play a strong part in AILI. GDF15 is a distinctive member of transforming growth factor beta (TGF-β) superfamily. It is a pleiotropic cytokine involved in regulating cell growth, anti-inflammation and anti-apoptosis (Unsicker et al., 2013). GDF15 can be rapidly induced by cellular stress and diverse disease conditions (Li et al., 2018a;Luan et al., 2019) including obesity (Mullican et al., 2017), cardiovascular disease (Wang et al., 2021a), cancer and nonalcoholic fatty liver disease (NAFLD). Previous reports suggested that GDF15 alleviated non-alcoholic liver steatohepatitis (NASH) in mice (Kim et al., 2018) and can be applied as a prognostic biomarker to predict advanced liver fibrosis in NAFLD patients (Koo et al., 2018). Growing evidence suggests a protective effect of GDF15 against chronic or acute liver injury in experimental animal models. GDF15 can reduce alcohol-induced fat accumulation and CCL4-induced fibrosis (Chung et al., 2017). Moreover, GDF15 inhibits LPS-induced acute liver injury by decreasing the activation of pro-inflammatory mediators (Li et al., 2018c). However, the precise role of GDF15 in AILI has yet to be explored.

In the present research, we investigated the temporal expression of Gdf15 in circulation and mouse liver tissues subject to APAP treatment. Overexpression of hepatic Gdf15 by AAV8-TBG-Gdf15 was used to investigate its therapeutic effect on AILI. Besides, recombinant human GDF15 (rhGDF15) were administered to explore its curative effect after APAP overdose. Unexpectedly, neither GDF15 overexpression nor recombinant protein of GDF15 could improve phenotype of APAP-induced hepatotoxicity.

## 2. Materials and methods

### 2.1 Chemicals and reagents

APAP were bought from MedChemExpress (cat. no. HY-66005/CS-2819). GSH Kit was purchased from NanJing JianCheng Bioengineering Institute (cat. no. A006-2-1). BCA Protein Assay Kits were obtained from Thermo Fisher Scientific (cat. no. 23227). rhGDF15was obtained from R&D Systems (cat. no. 957-GD-025/CF). Adeno-associated virus serotype 8 (AAV8) overexpressing Gdf15 was constructed and packaged by Weizhen Biosciences (Jinan, China). Antibodies against GDF15 (cat. no. sc-515675) was bought from Santa Cruz Biotechnology. Antibodies against p-JNK (cat. no. 4688T) was purchased from Cell Signaling Technology. Antibodies against JNK (cat. no. 66210-1-Ig) and tubulin (cat. no. 11224-1-AP) were bought from ProteinTech.

### 2.2 Experimental Animals

Male C57BL/6J mice (8-10 weeks old) were obtained from GemPharmatech Co., Ltd. (Nanjing, China). All mice were bred under specific pathogen-free conditions. All animal care and experiments were performed in accordance with the animal protocols and guidelines established by the Institutional Animal Use and Care Committee at the University of Science and Technology of China.

### 2.3 Animal treatments

#### 2.3.1 AAV8 overexpression

After randomization, mice were divided into three groups: (1) Normal saline + AAV8-GFP, (2) APAP + AAV8-GFP, (3) APAP + AAV8-GDF15. AAV8 was injected by tail vein injection (1 × 10^12^ vg per mice) twenty-one days before APAP treatment. Mice were injected with APAP (500 mg/kg) intraperitoneally after fasting overnight and sacrificed at 8 hours after APAP administration. Serum along with liver tissues were collected.

#### 2.3.2 Treatment with rhGDF15

Mice were randomly divided into three groups: (1) Normal saline, (2) APAP, (3) APAP + rhGDF15. Mice were performed by intraperitoneal (i.p.) injection of APAP (500 mg/kg) after overnight fasting. Two hours after APAP administration, normal saline or rhGDF15 (12 nmol/kg) was given by i.p.. Mice were sacrificed at 8 hours after APAP administration, and serum together with liver tissues were obtained.

### 2.4 Histological assessment

10% formaldehyde-fixed liver tissue embedded in paraffin. Then hematoxylin-eosin staining was performed and the pathological staining section was observed under light microscope.

### 2.5 Analysis of liver GSH amount

The content of GSH in liver was detected using GSH detection kit. Briefly, 30 mg liver tissues added with 270 μl normal saline were made into liver homogenate. The homogenate was centrifuged and the supernatant was extracted by reagents in accordance with the kit instruction. Absorbance was measured at 405 nm using microplate reader SpectraMax iD3 (Molecular Devices LLc, San Jose, USA). The corresponding protein levels of liver tissue was determined using BCA kit (Thermo Fisher Scientific, cat. no. 23227). Hepatic GSH levels were calculated using GSH content calibrated with protein concentration.

### 2.6 Quantitative real-time PCR analysis

The RNA of liver tissues (30 mg per sample) was extracted with tissue RNA purification kit plus (ES Science, Shanghai, China). Complementary DNA were synthesized by reverse transcription using PrimeScriptTMRT Master Mix (TaKaRa, Dalian, China). qPCR was achieved on a Roche LC96 Real-Time PCR Detection System with 2 × T5 Fast qPCR Mix (Tsingke Biotechnology, Beijing, China). The expression of detecting genes were calculated using the formula 2^−(ΔΔCt)^, and GAPDH serve as endogenous control. The primer sequences were shown in supplemental table 1.

### 2.7 Western blot analysis

Equal amount of protein from different liver samples were isolated by 10% SDS-PAGE gels and transferred to nitrocellulose membranes. Then, the membranes were blocked using Intercept® Blocking Buffer (LI-COR, 927-60001) and incubated overnight with corresponding antibodies at 4 °C. The primary antibodies included Gdf15, tubulin, p-JNK, and t-JNK. The immunoblot bands was visualized using LI-COR-CLx Infrared Imaging System (LI-COR, Lincoln, USA). Tubulin and t-JNK served as internal loading control for total protein and phosphorylated JNK, respectively.

### 2.8 GEO dataset from NCBI

One microarray dataset and two RNA-seq datasets from GSE17649 (mice liver after APAP treatment at 6 h and 0 h), GSE169155 (mice liver after APAP exposure at 6 h and saline treatment), and GSE111828 (mice liver of untreated control and APAP administration at 12 h) were obtained from GEO database (https://www.ncbi.nlm.nih.gov/gds). Differential expression genes (DEGs) were collected according to the selection criteria of *P* < 0.05 and fold change > 2. Venn diagram of differential expression genes intersection from the three datasets was characterized by Venny version 2.1 (https://bioinfogp.cnb.csic.es/tools/venny/).

### 2.9 Statistical analysis

Data are shown as mean ± standard deviation (SD). One-way ANOVA with Bonferroni post hoc test was used to make multiple comparisons. Statistical analysis was performed using GraphPad Prism 8. *P* < 0.05 was considered to be statistically different.

## 3. Results

### 3.1 Dynamic changes of hepatic GDF15 in response to APAP in mice

Gene profiles of GEO datasets including GSE17649, GSE169155, and GSE111828 were analyzed to identify DEGs after APAP overdose. GDF15 is one of the six common DEGs from the three datasets (Figure 1A). Remarkable increase expression of GDF15 was found at 6 h or 12 h after APAP administration in mouse liver. The specific parameters of GDF15 in the three datasets were presented in Figure 1B. Then, mice were given APAP (500 mg/kg) and collected liver tissue at 0 h, 3 h, 8 h, and 24 h (Figure 1C). As shown in Figure 1D, hepatocyte necrosis was induced and increased from 3 h to 24 h after APAP injection. Hepatocyte swelling and micro-alveolar changes were found at 3 h and 8 h, and a large area of vacuolar centrilobular necrosis was observed at 24 h. Hepatic protein expression of Gdf15 increased significantly at 3 h and 8 h after APAP overdose, and declined to normal level at 24h (Figure 1E).

**Figure 1.**
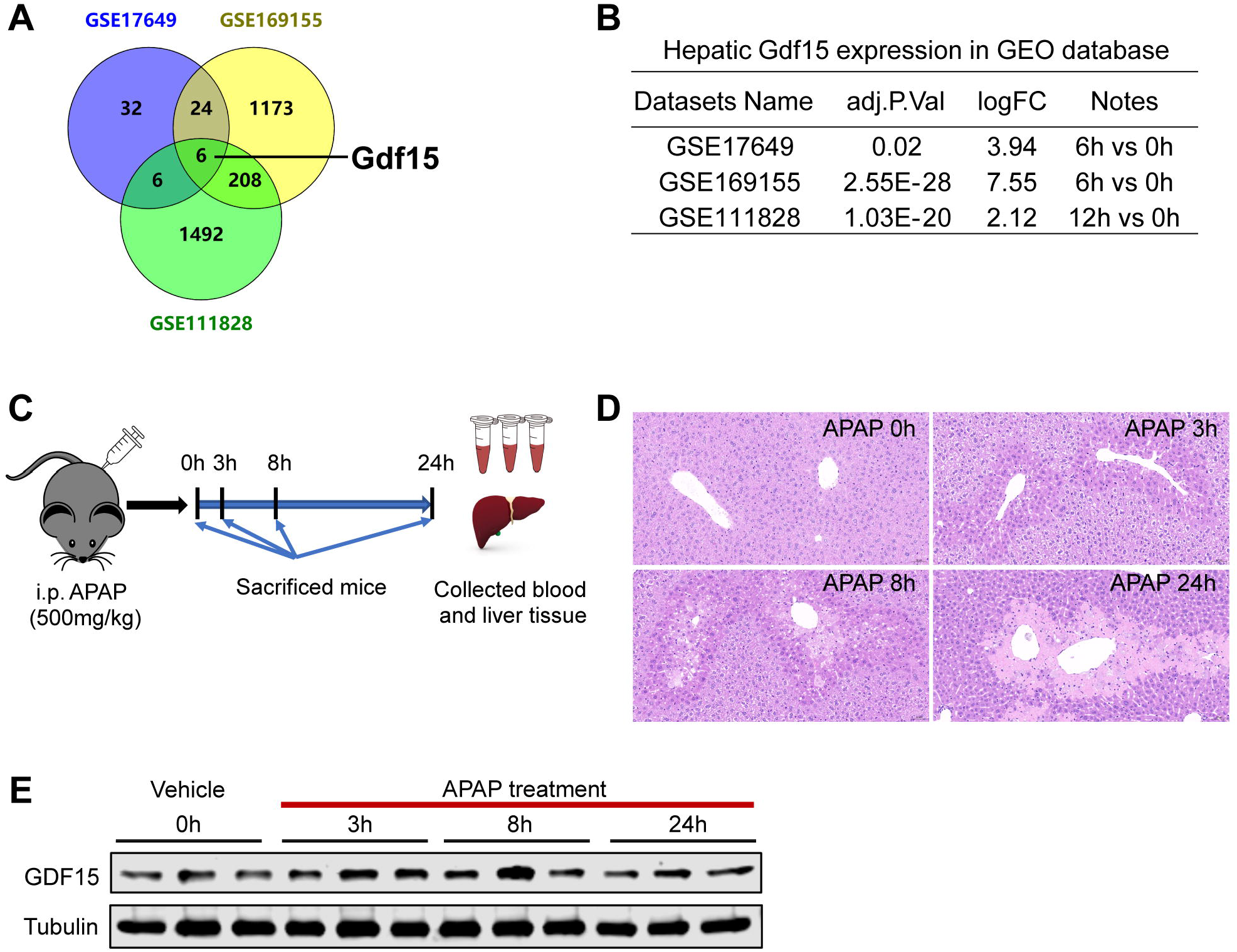
Temporal changes GDF15 expression after APAP treatment in mice. **(A)** Differential expression genes in Venn diagram constructed using three Gene Expression Omnibus (GEO) datasets including GSE17649, GSE169155, and GSE111828. **(B)** Hepatic expression of Gdf15 in GEO database. **(C)** Scheme of study design. **(D-E)** Male C57 BL/6J mice were given 500 mg/kg APAP by i.p. and sacrificed at 0 h, 3 h, 8 h, and 24 h after APAP administration (n = 5). **(D)** Representative H&E-stained (200 ×) liver sections at various time points. **(E)** Protein expression of hepatic Gdf15 at various time points.

### 3.2 Overexpression of hepatic GDF15 by AAV8 did not significantly change APAP induced hepatotoxicity

Considering its beneficial effects of GDF15 in many models of liver injury, overexpression of GDF15 using AAV8 was conducted to investigate its role in AILI (Figure 2A). Furthermore, we found that GDF15 was preferentially expressed in hepatocytes according to single-cell RNA sequencing results (https://tabula-muris.ds.czbiohub.org/) (Figure 2B). Therefore, TBG, a liver specific promoter, was added to AAV8 construction. As shown in Figure 2C, the mRNA levels of Gdf15 increased significantly at 8 h after APAP injection compared with normal saline group, and a marked elevation of Gdf15 in AAV8-Gdf15 group compared with AAV8-GFP group. Likewise, the protein expression of Gdf15 in AAV8-Gdf15 group was markedly increased than its expression in AAV8-GFP group (Supplemental Figure 1). Unexpectedly, there was no significant difference in serum ALT and AST between AAV8-Gdf15 group and AAV8-GFP group (Figure 2D-E). Besides, the area of necrosis (Figure 2F) and degree of apoptosis (Figure 2G) between the two groups did not change markedly. These results indicated that overexpression of hepatic Gdf15 did not affect liver injury induced by APAP.

**Figure 2.**
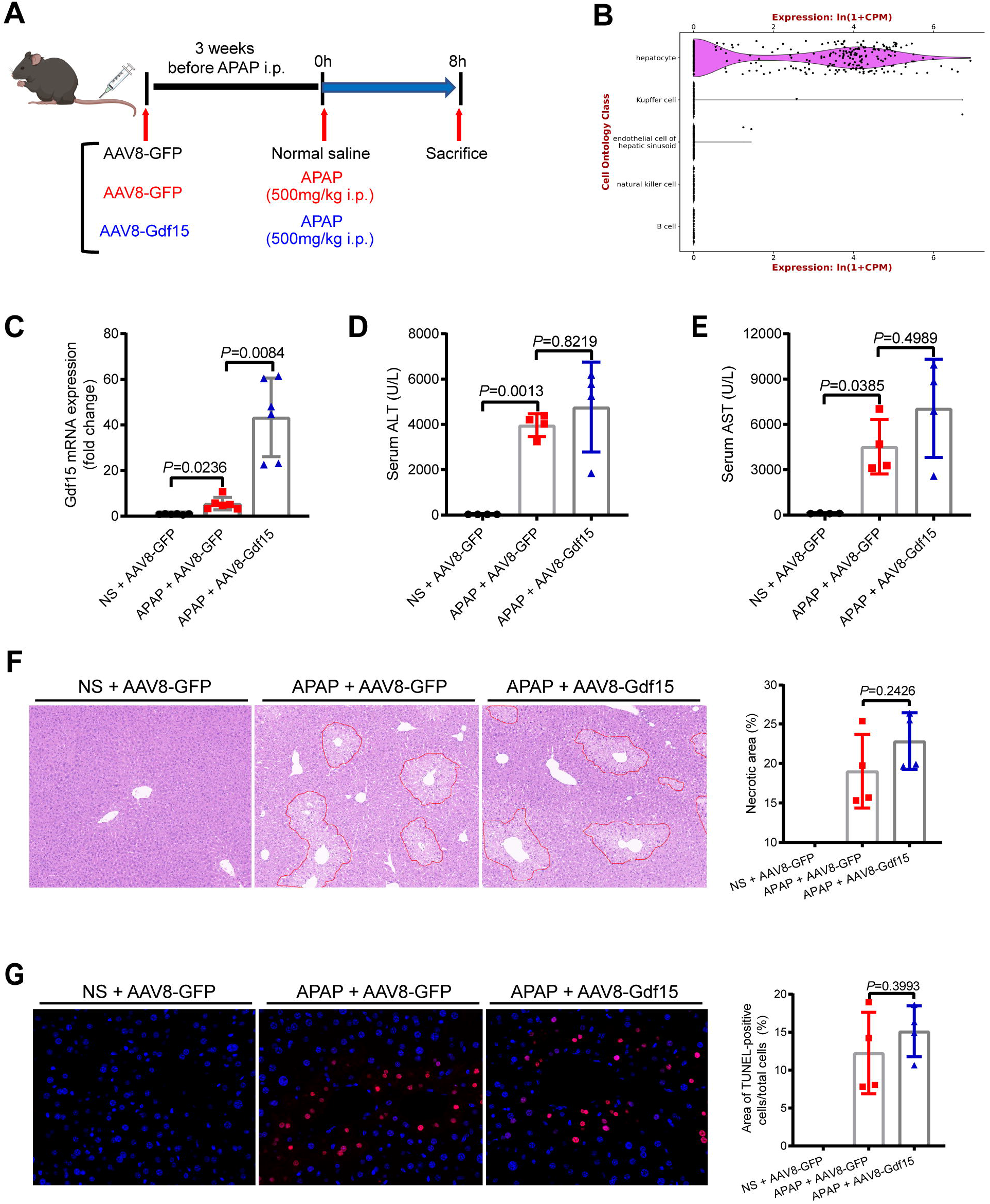
Overexpression of hepatic GDF15 had no significant effect on APAP-induced hepatotoxicity. **(A)** Schematic diagram of the present study. **(B)** Cell ontology class of Gdf15 in single-cell RNA sequencing data generated by the Tabula Muris consortium. **(C)** Hepatic mRNA expression levels of Gdf15 (n = 6). Serum contents of ALT **(D)** and AST **(E)** (n = 4). **(F)** Representative H&E-stained (100 ×) liver sections (left) and results of counted necrotic areas (right) (n = 4). **(G)** Representative TUNEL-stained (400 ×) liver sections (left) and area of TUNEL-positive cells/total cells (right) (n = 4). Data are represented as mean ± SD. *P* < 0.05 was considered as statistical difference.

To probe the molecular basis of Gdf15 overexpression in AILI, we measured the phosphorylation of JNK, hepatic GSH level, oxidative stress-related genes, and proliferation-associated genes. As a result, similar levels of p-JNK expression (Figure 3A-B), and glutathione content (Figure 3C) were detected in the livers of both AAV8-GFP and AAV8-Gdf15 mice treated with APAP. After APAP overdose, the expression of Gclc and Gclm increased significantly compared with normal saline group. Gdf15 overexpression exerted no noticeable impact on the expressions of antioxidative stress genes (Nrf2, Nqo1, Gclc, Gclm, and Sod2) (Figure 3D), and proliferation-associated genes (Pcna, Ccnb1, and Ccnd1) (Figure 3E).

**Figure 3.**
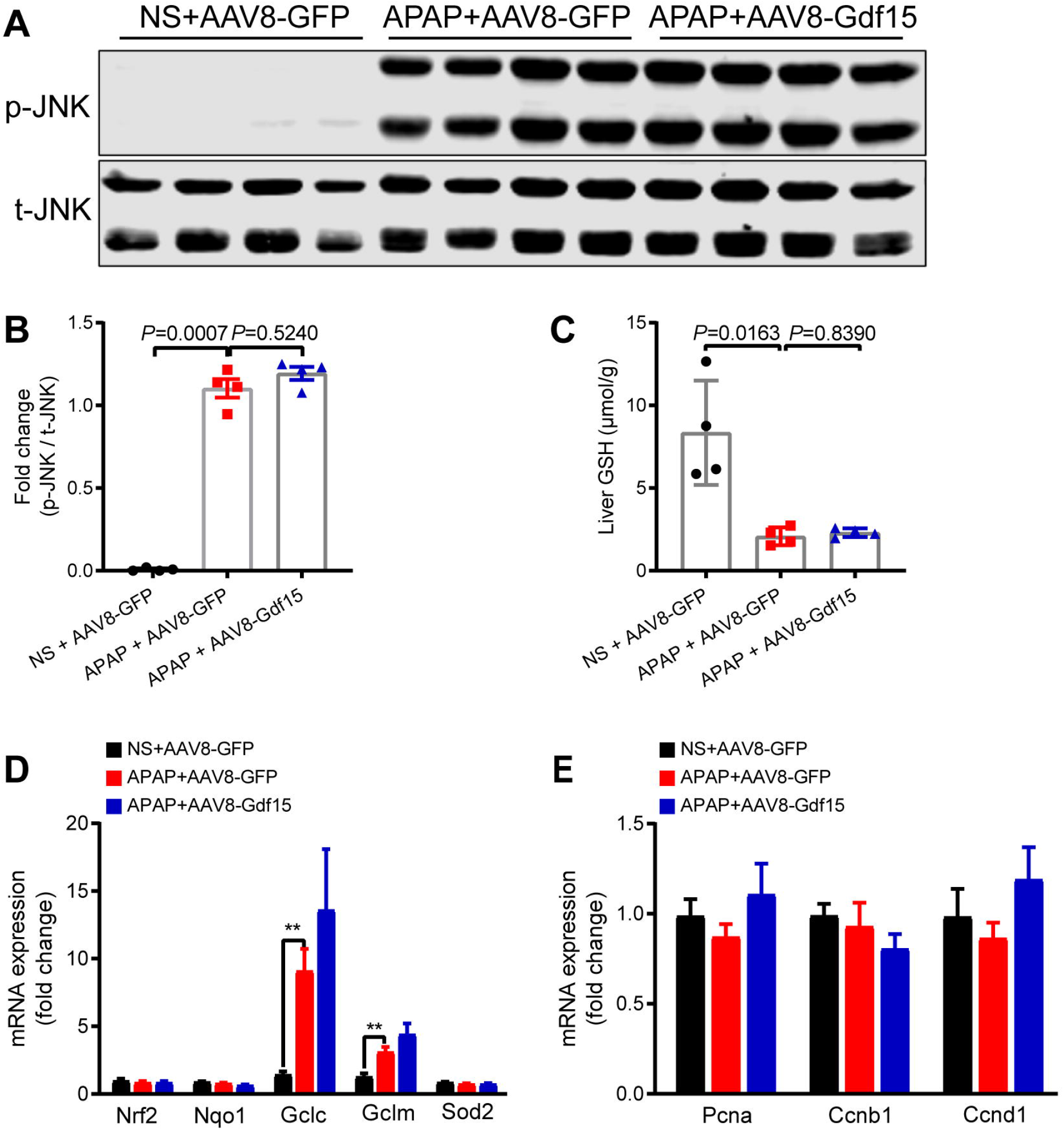
GDF15 overexpression did not significantly change hepatic p-JNK, glutathione levels, antioxidant genes, and proliferation-associated genes after APAP treatment in mice. **(A)** Protein expression of phosphorylated JNK (p-JNK) and **(B)** p-JNK protein expression quantified by densitometry (n = 4). **(C)** Glutathione (GSH) contents determined in liver tissues (n = 4). **(D)** Hepatic mRNA expression levels of antioxidative stress genes (Nrf2, Nqo1, Gclc, Gclm, and Sod2), and (E) antiproliferation-associated genes (Pcna, Ccnb1, and Ccnd1) (n = 6). Data are expressed as mean ± SD. \*\**P* < 0.01, \*\*\**P* < 0.001 versus normal saline group.

### 3.3 rhGDF15 administration did not improve liver injury induced by APAP

To further characterized the roles GDF15 in AILI, therapeutic administration of rhGDF15 (12nmol/kg body weight) was applied to investigate whether it can rescue AILI (Figure 4A). Similar to the results of GDF15 overexpression, serum ALT and AST (Figure 4B-C), hepatic area of necrosis (Figure 4D), and area of apoptosis (Figure 4E) did not markedly change after rhGDF15 administration.

**Figure 4.**
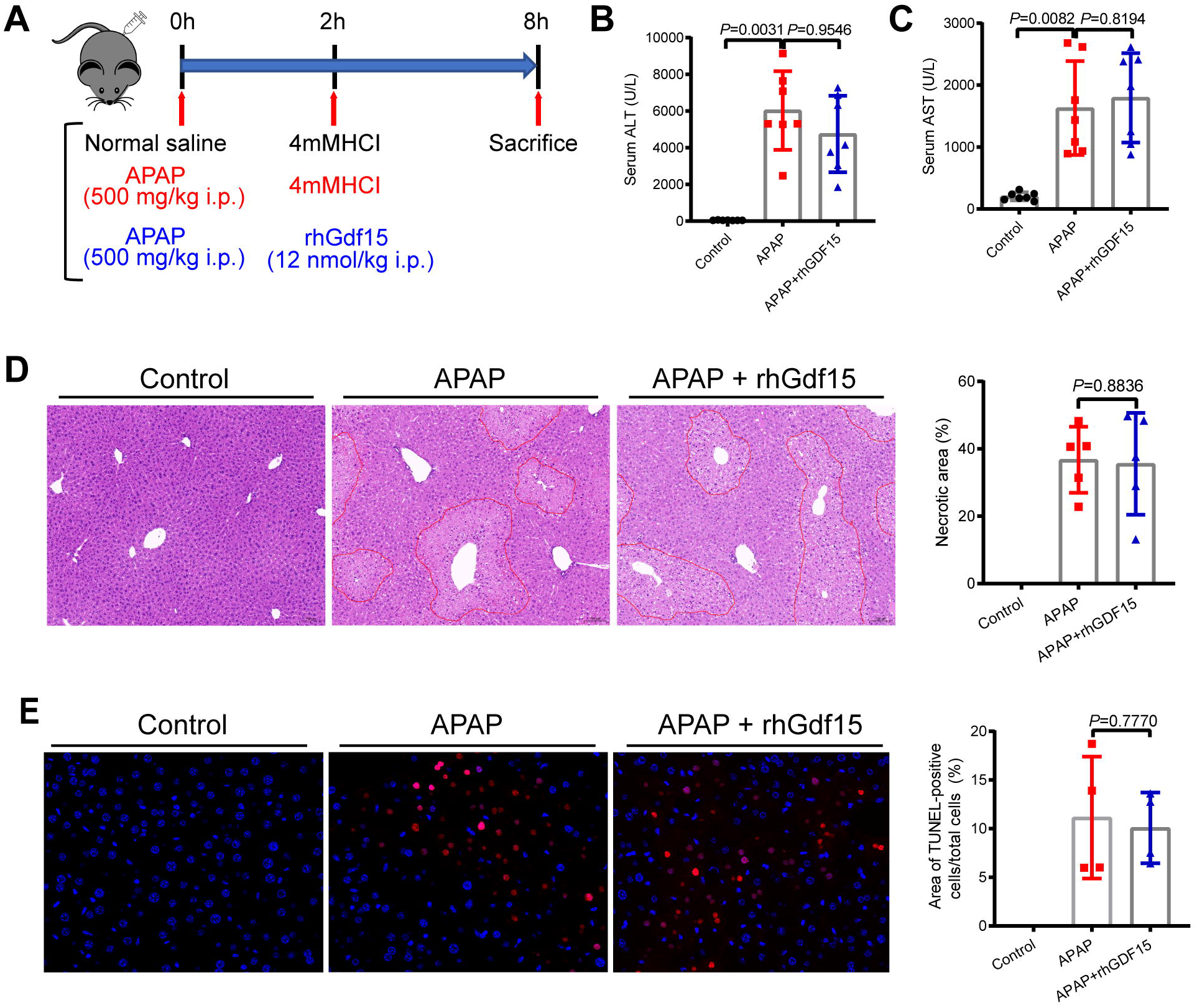
rhGDF15 administration did not change phenotype of APAP hepatotoxicity significantly. **(A)** Schematic diagram of the present study. Serum contents of ALT **(B)** and AST **(C)** (n = 7). **(D)** Representative H&E-stained (100 ×) liver sections (left) and results of counted necrotic areas (right) (n = 5). **(E)** Representative TUNEL-stained (400 ×) liver sections (left) and area of TUNEL-positive cells/total cells (right) (n = 4). Data are expressed as mean ± SD. *P* < 0.05 was considered as statistical difference.

Next, phosphorylation of JNK, hepatic GSH levels, oxidative stress-related genes, and proliferation-associated genes were detected between APAP group and rhGDF15 treatment group. As shown in Figure 5, the p-JNK protein level (Figure 5A-B), GSH level (Figure 5C), antioxidative stress genes (Nrf2, Nqo1, and Sod2) (Figure 5D), and proliferation-associated genes (Pcna, Ccnb1, and Ccnd1) (Figure 5E) did not change significantly between the two groups. Notably, rhGDF15 markedly increased the expression of Gclc and Gclm after APAP overdose (Figure 5D), suggesting rhGDF15 may enhance the antioxidant capacity despite the fact that rhGDF15 failed to alleviate APAP hepatotoxicity.

**Figure 5.**
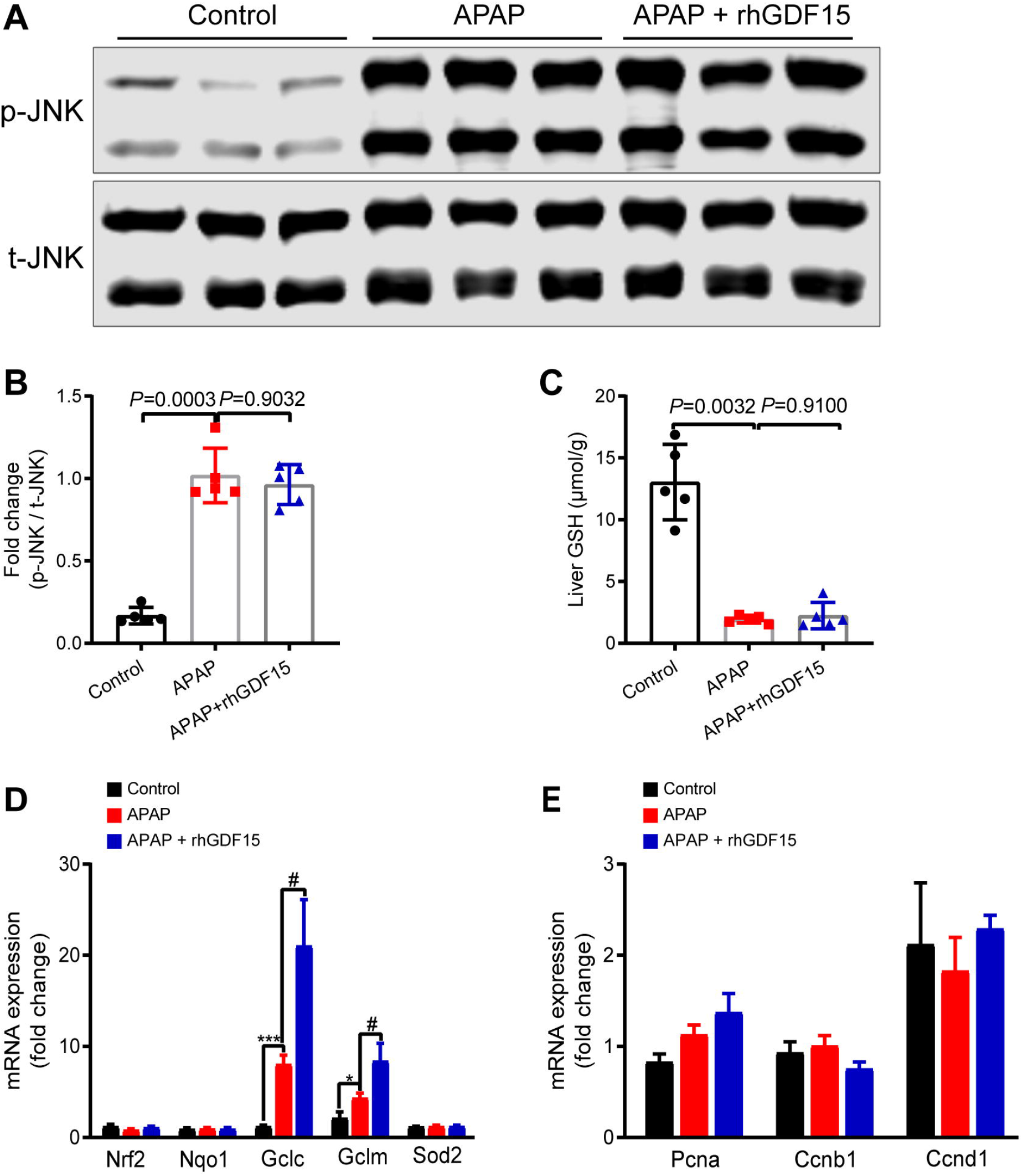
rhGDF15 administration did not significantly change hepatic p-JNK, glutathione level, antioxidant genes, and proliferation-associated genes after APAP treatment in mice. **(A)** Protein expression of phosphorylated JNK (p-JNK) and **(B)** quantified by densitometry (n = 5). **(C)** Glutathione (GSH) contents determined in liver tissues (n = 5). **(D)** Hepatic mRNA expression levels of antioxidative stress genes (Nrf2, Nqo1, Gclc, Gclm, and Sod2), and **(E)** antiproliferation-associated genes (Pcna, Ccnb1, and Ccnd1) (n = 6). Data are expressed as mean ± SD. \**P* < 0.05, \*\*\**P* < 0.001 versus normal saline group. ^#^*P* < 0.05 versus APAP group.

## 4. Discussion

In this research, we found increased expression of GDF15 after APAP treatment at different timepoint in mice. Drastic elevation of GDF15 was also presented in GEO database at 6 h or 12 h after APAP overdose. Previous studies indicated that GDF15 could alleviate NASH in mice (Kim et al., 2018) or inhibited LPS-induced acute liver injury with anti-inflammatory effects (Li et al., 2018c), suggesting a beneficial effect of GDF15 on liver disease or liver injury in experimental models. Based on the hepatoprotective effects of GDF15 and the fact that therapeutic approach of AILI is limited, we hypothesized that GDF15 may ameliorate APAP hepatotoxicity. However, overexpression of hepatic GDF15 or therapeutic administration of rhGDF15 was not able to improve phenotype of APAP hepatotoxicity significantly. Similarly, one previous research proved no difference in liver injury in GDF15 deficient mice compared with wild-type mice in partial-hepatectomy and carbon tetrachloride experimental models (Hsiao et al., 2000). It is possible that the increased expression of GDF15 induced by cell stress or tissue injury may be a compensatory survival response with no regulatory action. In present study, we found no obvious therapeutic effect of GDF15 on AILI under our experimental conditions. This does not exclude that knockdown GDF15 or prevention administration of rhGDF15 will affect AILI.

Nrf2 is an essential factor of oxidative stress which drives the expression of cytoprotective genes, including GCLC, GCLM, heme oxygenase 1 (HO-1), and superoxide dismutase 2 (SOD2) (Zhang et al., 2017). Nrf2 exerts a crucial role in AILI (Chan et al., 2001). In this study, the expression of Nfr2, HO-1, and SOD2 did not change significantly. A noticeable elevation of GCLC and GCLM was observed after rhGDF15 administration. Moreover, GCLC and GCLM are vital enzymes which catalyze the synthesis of glutathione (Lu, 2013). Increase of GCLC and GCLM may contribute to increased synthesis of glutathione. Glutathione plays a pivotal role in antioxidant and detoxication in AILI (Jaeschke et al., 2020). At 8 h after APAP administration, level of glutathione was depleted for binding with toxic metabolites of APAP. There was no remarkable difference in glutathione content between control group and GDF15 overexpression group. Although rhGDF15 induced the expression of GCLC and GCLM, which may boost antioxidative stress ability and increase glutathione synthesis after APAP overdose, this protection was limited compared to drastic depletion of GSH in AILI. This mechanism may underline the inability of GDF15 to alleviate AILI.

GDF15 is a stress and disease-induced cytokine which can be secreted into the circulation. Circulating GDF15 may serve as a promising biomarker of aging (Liu et al., 2021), cardiovascular disease and mortality associated with all outcomes (Ho et al., 2018). GDF15 can also predict the adverse outcomes of high-risk obese patients (Sarkar et al., 2020). Recently, GDF15 receives intensive research interest in obesity treatment for the identification of its orphan receptor, glial-derived neurotrophic factor (GDNF) receptor alpha-like (GFRAL) (Emmerson et al., 2017;Hsu et al., 2017). Numerous drug development and mechanistic research in the treatment of metabolic disorders are being conducted on GDF15/GFRAL signaling pathway which provides new insights into body weight control (Coll et al., 2020;Wang et al., 2021a). Nevertheless, some key questions remain in the field of GDF15. Tissue source or derivation from which type of cells remain largely elusive. Under physiological conditions, high expression of GDF15 is observed in kidney, liver, adipose tissue, and skeletal muscle (Patel et al., 2019). Under pathological condition such as obesity or inflammation stimuli, elevation of GDF15 from immune cells can infiltrate liver and this may result in increase of GDF15 (Liu and Nikolajczyk, 2019). Single-cell RNA sequencing of human liver specimens from healthy or NAFLD revealed increased GDF15 expression in almost all types of liver cells (Ramachandran et al., 2019;Govaere et al., 2020). Whether GDF15 from liver contributes to its increase of circulatory levels warrants further studies. Secretory GDF15 in circulation is a vital biomarker induced in many diseases and tissue damage. The synthesis of GDF15 in circulation exhibits diverse forms such as pre-pro-GDF15, pro-GDF15, and mature GDF15 (Li et al., 2018b). The levels and function of its precursor or mature protein either in monomer or homodimer form are still largely unknown. Of note, GDF15 exerts a vital part in obesity, NAFLD and other diseases. The mechanism for controlling body weight depends on GDF15/GFRAL signaling pathway (Yang et al., 2017). However, the expression of GFRAL was only found in the area postrema and nucleus of the solitary tract of hindbrain (Hsu et al., 2017). It remains to identify whether GDF15 is protective against other types of liver injury and the putative receptors involved.

In summary, GDF15 expression was significantly elevated upon APAP overdose. However, overexpression of GDF15 or administration of rhGDF15 was incapable to ameliorate APAP-induced hepatotoxicity Despite significant increase of Gclc and Gclm gene expression in rhGDF15-treated mouse liver. Despite existing evidence suggesting the hepatoprotective role of GDF15, GDF15 fails to ameliorate liver injury induced by APAP.

## Data availability statement

All the data listed in this article will be available to any qualified researcher.

## Author contributions

SX and JW conceived and designed the experiments. PJ and ZL performed the experiments and wrote the paper. PJ, ZL and SX collected and analyzed the data. TF and ZZ assist with experiments. DW provide insightful comments on the manuscript and proof-read the manuscript. All authors have read and approved the final manuscript.

