## Supplemental material for "Effect of GDF15 on acetaminophen (APAP)-induced liver injury in mice"

### Supplementary Data

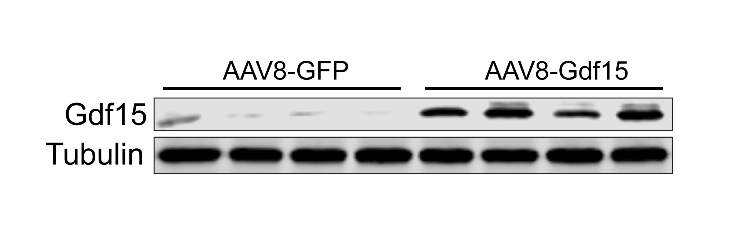

**Supplementary Figure 1.** The protein expression of Gdf15 between AAV8-GFP group and AAV8-Gdf15 group.

**Supplementary Table 1.** Sequences of primers used for real-time PCR

| Primer name | Sequence from 5' to 3' |
| --- | --- |
| Nrf2 forward | TAGATGACCATGAGTCGCTTGC |
| Nrf2 reverse | GCCAAACTTGCTCCATGTCC |
| Nqo1 forward | AGGATGGGAGGTACTCGAATC |
| Nqo1 reverse | TGCTAGAGATGACTCGGAAGG |
| Gclc forward | GGACAAACCCCAACCATCC |
| Gclc reverse | GTTGAACTCAGACATCGTTCCT |
| Gclm forward | CTTCGCCTCCGATTGAAGATG |
| Gclm reverse | AAAGGCAGTCAAATCTGGTGG |
| Sod2 forward | CAGACCTGCCTTACGACTATGG |
| Sod2 reverse | CTCGGTGGCGTTGAGATTGTT |
| Pcna forward | TTTGAGGCACGCCTGATCC |
| Pcna reverse | GGAGACGTGAGACGAGTCCAT |
| Ccnb1 forward | AAGGTGCCTGTGTGTGAACC |
| Ccnb1 reverse | GTCAGCCCCATCATCTGCG |
| Ccnd1 forward | GCGTACCCTGACACCAATCTC |
| Ccnd1 reverse | CTCCTCTTCGCACTTCTGCTC |
| Gdf15 forward | ACGCATGCGCAGATCAA |
| Gdf15 reverse | CACTGTCTGTCCTGTGCATAA |
| Gapdh forward | AACAGCAACTCCCACTCTTC |
| Gapdh reverse | CCTGTTGCTGTAGCCGTATT |
